# Mutational Analysis of SARS-CoV-2 Genome in African Population

**DOI:** 10.1101/2020.09.07.286088

**Authors:** Olabode E. Omotoso, Ayoade D. Babalola, Amira Matareek

## Abstract

Severe acute respiratory syndrome coronavirus 2 (SARS-CoV-2), a highly infectious and pathogenic virus has claimed lot of lives globally since its outbreak in December 2019 posing dire threat on public health, global economy, social and human interaction. At moderate rate, mutations in the SARS-CoV-2 genome are evolving which might have contributed to viral genome variability, transmission, replication efficiency and virulence in different regions of the world. The present study elucidated the mutational landscape in SARS-CoV-2 genome among the African population, which may have contributed to the virulence, pathogenicity and transmission observed in the region. Multiple sequence alignment of the SARS-CoV-2 genome (356 viral protein sequences) was performed using ClustalX version 2.1 and phylogenetic tree was built using Molecular Evolutionary Genetics Analysis (MEGA) X software. ORF1ab polyprotein, spike glycoprotein, ORF3, ORF8 and nucleocapsid phosphoprotein were observed as mutational hotspots in the African population and may be of keen interest in the adaptability of SARS-CoV-2 to the human host. While, there is conservation in the envelope protein, membrane glycoprotein, ORF6, ORF7a, ORF7b and ORF10. The accumulation of moderate mutations (though slowly) in the SARS-CoV-2 genome as revealed in our study, could be a promising strategy to develop drugs or vaccines with respect to the viral conserved domains and host cellular proteins and/or receptors involved in viral invasion and replication to avoid a new viral wave due to drug resistance and vaccine evasion.

## Introduction

The novel coronavirus caused by severe acute respiratory syndrome coronavirus 2 (SARS-CoV-2) was first reported in December 2019 in Wuhan, China. The virus was initially named: Wuhan seafood market pneumonia virus before the disease was officially named coronavirus disease 2019 (COVID-19) by World Health Organization (WHO) on February 12, 2020. As at September 5, 2020, 08:42 CEST COVID-19 has spread in 216 countries with 26,994,442 confirmed cases and 880,994 deaths, which accounts for 1,293,048 confirmed cases and 31,082 deaths in Africa [1], [2]. There exists a vast difference in the fatality rates across countries; this might be a result of varied demographic structure, adherence to public health guidelines (wearing of face masks, frequent hand washing under running water with soap and use of alcohol-based sanitizers) and the measures (physical and social distancing, quarantine, self-isolation, partial or full-scale lockdown) put in place in different countries to curtail viral transmission. At the early stage of infection, infected persons could be asymptomatic while some experience mild to moderate symptoms, such as high fever, cough, and headache and, in severe cases, pneumonia [3]. Infected persons with underlying health conditions such as chronic lung disease, diabetes or hypertension are at high risk for severe symptoms and eventual death from the disease.

Coronaviruses are large, enveloped, single-stranded RNA viruses with the largest genome (ranging from 26 to 32 kb) among all RNA viruses. The viral genome structure consists of a 3□-, 5□-untranslated regions (UTRs) encoding ORF1ab polyproteins, ORF3a, ORF6, ORF7a, ORF7b, ORF8, surface or spike glycoprotein (S), envelope (E), membrane (M), and nucleocapsid (N) proteins [4], [5]. The Global Initiative on Sharing All Influenza Data (GISAID) have identified three major clades of the virus, named clade G (spike glycoprotein variant - D614G), clade V (ORF3a variant - G251V), and clade S (ORF8 variant - L84S). Although earlier report [6] have confirmed a low viral genomic variability, the question still remains if viral transmission, incidence and fatality observed in different regions could be attributed to geographic distributions of viral clades and variants.

Comprehensive genomic analysis of epidemiologic viral sequence coming from different countries in a region can help in gaining insights into the SARS-CoV-2 virulence and pathogenesis. To date, no precise antiviral vaccine or therapy has proven effective. Thus, genomic variability studies of the virus can play an important role in gaining insight to the genomic diversity in order to adopt measures to curtail the menace of COVID-19. Scientific findings and complete viral genome sequences have been made publicly available, thanks to researchers, healthcare workers, the GISAID consortium and National Center for Biotechnology Information (NCBI), among many others. This data avalanche will aid rapid effort in understanding viral genome diversity, and identification of suitable targets for drugs and vaccine development [7]. Biological description of viral RNA mutations can provide information for assessing viral transmission, drug resistance, host invasion and pathogenesis [8]. In the present study, we investigated the extent of whole genome variation in African population. We stratified recurrent mutations, and also identified the mutational hotspots and conserved domains in the viral genome. These findings may help to understand the pattern of viral transmissibility, evolution and virulence in Africa, as well as identifying possible targets for antiviral drugs, vaccines, diagnostic assays and improved control of the pandemic.

## Materials and Methods

### Data acquisition

Whole-genome protein sequences of SARS-CoV-2 isolates were downloaded from GISAID (https://epicov.org/epi3) and NCBI database (https://ncbi.nim.nih.gov/genbank/sars-cov-2-seqs/). A total of 356 SARS-CoV-2 complete protein sequences (Northern Africa - 150, Western Africa - 102, Eastern Africa – 54, South Africa - 41, and Central Africa – 9) were mined, after filtering as complete, high coverage only, low coverage excluded, and of Africa descent. The regions infected by SARS-CoV-2 and that shared the complete SARS-CoV-2 genomes used in this study are provided as Supplementary file S1.

### Multiple sequence alignment (MSA) and phylogenetic tree

The mined SARS-CoV-2 protein sequences were aligned with the reference genome (accession number: NC_045512.2) by using ClustalX version 2.1 [9] MSA software and the default parameters [10]. The evolutionary history of the viral sequences was inferred by using the Maximum Likelihood method and Jones-Taylor-Thornton (JTT) matrix-based model [11]. The tree with the highest log likelihood (−21700.94) is shown. Initial tree(s) for the heuristic search were obtained automatically by applying Neighbor-Join (NJ) and BioNJ algorithms to a matrix of pairwise distances estimated using the JTT model, and then selecting the topology with superior log likelihood value. The tree is drawn to scale, with branch lengths measured in the number of substitutions per site. Evolutionary analyses were conducted in Molecular Evolutionary Genetics Analysis (MEGA) X software [12].

### Sequence and mutational analysis

In this study, we used the mined data to analyze the variability in the SARS-CoV-2 genome in the African population since the onset of the pandemic in order to identify its conserved domains and mutational patterns. Recurrent mutations were focused on as they are likely candidates for the ongoing adaptive strategies of SARS-CoV-2 to human host and possible target for drug development, while sites with undetermined residues (labeled as X), few or single variation occurrence were excluded.

## Results

Multiple sequence alignment of the 356 SARS-CoV-2 protein sequences revealed high rate recurrent mutations at 23 distinct sites in the ORF1ab polyprotein sequence. Very high rate recurrent mutations exist at the following sites: nsp2 region; T265I with occurrence in 24 viral sequence, nsp3 region; D1037E which occurred in 11 viral sequence, T1246I in 9 viral sequence, and E1363D in 11 viral sequence, nsp4 region; E3073A in 9 viral sequence, 3 chymotrypsin-like proteinase region (3CL^pro^) G3278S in 11 viral sequence, nsp6; L3606F in 21 viral sequence, nsp8; T4090I in 10 viral sequence, and RNA dependent RNA polymerase (RdRp) region; P4715L. The P4715L mutation occurred in 280 sequences (78.7%) flagging the RdRp region as a mutational hotspot. The D1037E, G3278S and T4090I mutations concurrently occurred in 10 viral sequences all from Egyptian population. Among the nsp’s, RdRp has more variants (288) in the analyzed viral sequences, followed by nsp3 and nsp6 with 49 and 21 mutated viral sequence respectively. These recurrent mutations characterized the parsim information responsible for clusters in the phylogenetic tree (Suppl. file S2).

In the spike glycoprotein region, recurrent mutations were observed in the S1 domain; R408I in 4 viral sequence, and S2 domain of the spike glycoprotein; Q677H in 9 viral sequences respectively. The highly recurrent D614G mutation in the S1 domain was observed in 288 viral sequences (80.9%), flagging this position as a SARS-CoV-2 mutational hotspot. The D614G mutation was observed as follows; Northern Africa (43%), Western Africa (20.5%), Southern Africa (17%), Eastern Africa (16.3%), and Central Africa (3.2%).

Q57H recurrent mutation in the ORF3 protein region was observed in 104 viral sequence (Northern Africa – 52%, West Africa – 38%, Eastern Africa – 8%, Central Africa – 2% and no occurrence in Southern African sequences) and S171L variant in 6 sequences (Egypt – 4, and Mali - 2). The L84S mutation in the ORF8 region was observed in 41 viral sequences. The L84S mutation was predominant in West Africa (65.85%), Northern Africa (19.51%) and Eastern Africa (12.2%), with very few/no observance in viral sequences from Central and Southern Africa. Recurrent mutations in the viral genome are shown in Table 1 and Figure 1.

**Table 1.** Overall mutation distribution in the SARS-CoV-2 genome in African population

| **Viral genome** | **Number of sites mutated** | **Frequency of mutations** |
| --- | --- | --- |
| ORF1ab/ORF1a | 25 | 468 |
| Spike glycoprotein | 5 | 305 |
| ORF3a | 3 | 114 |
| Envelope protein | 1 | 2 |
| Membrane glycoprotein | 1 | 1 |
| ORF6 | 0 | 0 |
| ORF7a | 1 | 2 |
| ORF7b | 0 | 0 |
| ORF8 | 2 | 43 |
| Nucleocapsid phosphoprotein | 6 | 195 |
| ORF10 | 0 | 0 |

\*Highly recurrent mutations were observed in ORF1ab polyprotein, Spike protein, ORF3a, and Nucleocapsid phosphoprotein regions, flagging these regions as mutational hotspots

**Figure 1.**
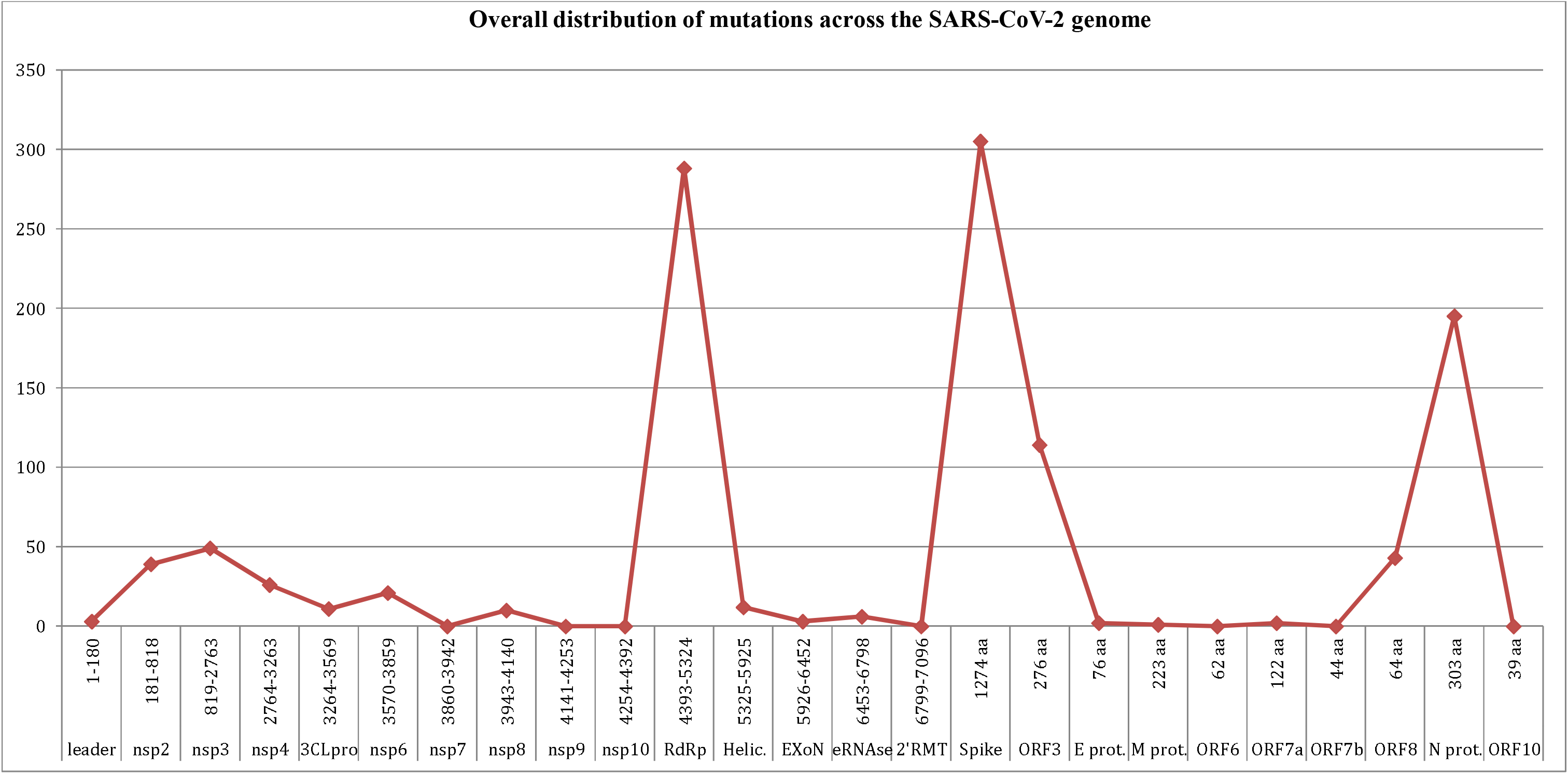
Distribution of mutations in the 356 SARS-CoV-2 protein sequences. We observed very high rate recurrent mutations in multiple sequences at the RNA-dependent RNA polymerase (RdRp) coding region, spike glycoprotein, ORF3, nucleocapsid phosphoprotein region.

Concurrent recurrent mutations were observed in the Nucleocapsid phosphoprotein; R203K and G204R in 74 viral sequences, S202N variant was observed in 35 viral sequences (West Africa – 76%, East Africa – 17%, and North Africa – 7%) and S187L in 8 sequences. The R203K and G204R mutations were prevalent in South Africa (40.54%), West Africa (32.43%), and East Africa (17.6%) and with little/no occurrence in viral sequences from Central African and Northern African countries. The S187L mutations in this study were all observed in Senegalese population.

## Discussion

The genome is the molecular architecture of any life, dictating its genotypic and phenotypic expression. Although at a minimal rate, mutations in the SARS-CoV-2 genome are evolving which might have contributed to viral genome variability, transmission, replication efficiency and virulence in different regions of the world [13]. We performed mutational analysis of the SARS-CoV-2 genome to understand the epidemiology, extent of spread and implication of mutations in the African population and also to identify loopholes in the viral genome as potential target for drug or vaccine development. In the African population, the SARS-CoV-2 ORF1ab polyprotein, spike glycoprotein, ORF3, ORF8 and nucleocapsid were observed as mutational hotspots due to high rate of recurrent mutations. While, there is conservation in the envelope protein, membrane glycoprotein, ORF6, ORF7a, ORF7b and ORF10.

ORF1ab polyprotein, which is the longest ORF in the SARS-CoV-2 genome, is cleaved into many nonstructural proteins (nsp1-nsp16). As at August 14, 2020, the P4715L mutation has been observed in 60973 viral sequences (76.94%) in 97 countries and the T265I mutations 13765 times (17.39%) in 72 countries (https://epicov.org/epi3), while D614G spike mutation has been observed in 61298 viral sequences (77.23%) in 96 countries and Q57H mutation 17727 times (22.38%) in 87 countries (https://epicov.org/epi3). L84S mutation at ORF8 has been observed in 5686 viral sequences (7.18%) in 62 countries, while the concurrent R203K and G204R mutations have occurred 25503 and 25430 times (32%) respectively in 83 countries (https://epicov.org/epi3).

The ORF1ab polyprotein is essential for genome replication. RdRp is responsible for viral RNA replication, thus, it is expected that RdRp is well conserved. Surprisingly, our study in line with earlier findings [8], [13] has reported recurrent mutations in this region resulting in alteration of protein sequence. In particular, the P4715L mutation (present in 280 viral sequences (78.7%) in our study) located close to a hydrophobic cleft has been identified as a potential target for antiviral drugs [8]. The use of molecular docking has identified Atazanavir as a potential candidate for COVID 19 drug due to its binding affinity to RdRp (Kd 21.83 nM) [14]. Interestingly, most of the viral sequences bearing the P4715L mutation also concurrently bear the D614G mutation in the Spike glycoprotein; this could suggest a synergistic effect of these mutations in these 2 important domains.

The SARS-CoV-2 3CLpro enzyme controls viral replication and is vital for its life cycle. During the disease outbreak of SARS-CoV and Middle East respiratory syndrome coronavirus (MERS-CoV), 3CLpro was a proven drug discovery target. Molecular docking has also identified potential candidates as antiviral drugs; aliskiren, dipyridamole, mopidamol and rosuvastatin with a relatively high binding energy to viral 3CLpro [14]. In our study, G3278S mutation was only observed in this region in 9 viral sequences, which were all Egyptians. Nevertheless, the importance and the relative conservativeness of this region could make it a suitable target for antiviral inhibitory screening, as suggested in earlier study [15].

The spike (S) glycoprotein mediates host cell-surface receptor binding via its S1 domain and induces fusion of host and viral membranes through the S2 domain and determines viral transmission, invasion and host tropism [16]. These are the initial and critical steps in the coronavirus infection cycle; this can also serve as potential targets for antiviral drug design. The spike glycoprotein which exists in the prefusion or postfusion conformations [16] binds to the human angiotensin-converting enzyme (ACE) 2 receptor to gain cellular entry, which facilitates its rapid spread to all parts of the globe. Hence, mutations in the S glycoprotein may result to decreased binding affinity for the ACE2 protein, thereby affecting viral fitness. The S glycoprotein being a surface protein is constantly under selective pressure in a bid to evade immune response; this might have promoted its adaptation to the host genome and the recurrent mutation observed. Earlier report before the outbreak of SARS-CoV-2 has shown that mutations in the human SARS-CoV spike S1 domain allows it to bind much more tightly to human ACE2 compared to civet SARS-CoV S1 [16]. We observed mutation hotspots (D614G) in the S1 domain with a relatively conserved S2 domain, which may infer membrane fusion as the central function of SARS-CoV-2 S glycoprotein. Previous study has likewise established that coronaviruses can elicit receptor-independent viral entry into host cells [16]. Therefore, more focus should be given to understanding the mechanism of S2 domain mediating host-cell membrane fusion as potential cellular targets for antiviral therapy [14].

Nucleocapsid phosphoprotein (N) composed of the C- and N-terminal domain forms the ribonucleoprotein complex with the viral RNA; it enhances the viral genome transcription and facilitates the interaction with the membrane protein during virion assemblage [17]. Mutations in the viral nucleocapsid region alters its binding to miRNAs, which might play a role in the pathogenesis and progression of infection in the patient [13]. Save for 15 viral sequences from Kenya, the occurrences of P4715L and L84S (28144T>C) coincide, this corroborates earlier study where L84S in the viral ORF8 is always found in pair with a synonymous mutation, 8782C>T [3].

The membrane glycoprotein (M) determines the shape of the viral envelope and the envelope protein (E) plays crucial role in viral assembly and budding [17]. The M protein (~25–30 kDa) consist three transmembrane domains. After viral membrane fusion, the interaction of S with M protein is necessary to retain S in the Endoplasmic Reticulum-Golgi intermediate compartment (ERGIC)/Golgi complex and its integration into new virions [17]. The M protein also binds to N protein in order to stabilize the nucleocapsid and aid viral assembly. During viral replication, E protein is upregulated in the infected cell facilitating viral assembly. The role of E protein in viral maturation have been expressed in E protein knock-out recombinant coronaviruses, with resultant crippled viral maturation, and reduced viral titres [17]. Our findings revealed a well conserved E and M protein, thus based on the importance of these structural proteins; they can serve as potential antiviral therapeutic target; as earlier attempted [18] by altering recombination causing large deletions in the E proteins.

The viral ORF3a and ORF10 proteins can synergistically attack heme on the host’s hemoglobin 1-β chain, thereby disintegrating iron to form porphyrin, which will therefore result to less and less hemoglobin carrying oxygen and carbon dioxide, interfering with the heme pathway and extreme poisoning and inflammation of the hepatocytes [14]. Studies on chloroquine (CQ) and hydroxychloroquine (HCQ) have shown the inhibitory activities against viral S protein and ORF8 binding to porphyrin and against the viral ORF1ab, ORF3, and ORF10 proteins attacking heme to form porphyrin, thus easing respiratory distress symptoms [14]. The use of CQ and HCQ as a potent drug against coronavirus has generated a lot of controversies due to their adverse effects in patients. Meanwhile, on Jun 16, 2020, the US Food and Drug Administration (FDA) retracted the use of CQ and HCQ as a potent candidate for coronavirus treatment due to their lack of efficacy and safety concerns (http://www.chinadaily.com.cn/). Hence, the quest for a clinically approved and efficient therapeutic agent is still on and our study has been able to suggest potential targets for drugs or vaccine development.

Due to our keen interest in mutations that have effect on protein sequence, synonymous mutations which do not alter amino acid residue were not accounted for in the present study. More so, this genomic dataset includes data from single country each from Central Africa and South Africa, therefore, some of the genomic diversity might likely remain unsampled. Hence, we encourage support for biomedical researchers and research institutes in developing countries in order to generate extensive genomic resources to understand viral transmissibility, evolution and variation in the African region.

## Conclusion and perspective

Conclusively, SARS-CoV-2 is a highly infectious and pathogenic virus that has claimed hundreds of thousands of lives globally since outbreak in December 2019. Our study has elucidated the mutational landscape and conserved domains in the SARS-CoV-2 genome in the African population, which may have contributed to its virulence, pathogenicity and transmission in the region. Due to the accumulation of moderate mutations (though slowly) in the SARS-CoV-2 genome as revealed in our study, drugs targeting the viral proteins might not be as efficient as those designed to target host’s cellular proteins and/or receptors. Hence, it will be a promising strategy to develop drugs or vaccines with respect to the viral conserved domains and host proteins involved in viral invasion and replication to avoid a new viral wave due to drug resistance and vaccine evasion. Africa is blessed with traditional plants that have been used over the years in managing and treating wide spectrum of diseases, hence, combination of traditional African medicine and other candidate antiviral drugs might have better therapeutic prospect. No doubt, the traditional plants and candidate medicines will require extensive clinical trials to ascertain the safety concerns, mechanism of action, adverse effects, and efficacy.

## CRediT authorship contribution statement

OO: Conceptualization, Data mining, Methodology, Analysis, Writing - original draft. DB: Data sorting, Methodology, Analysis, Writing - review & editing. AM: Data mining, Analysis, Writing – review and editing.

## Declaration of Competing Interest

All authors declare no conflict of interest

## Funding

No funding received for this study

## Acknowledgements

The authors would like to appreciate the heroic effort of health-workers in the frontline of tackling COVID19 and all researchers who sequenced SARS-CoV-2 genomes and deposited them in public repositories (NCBI and GISAID) used in this study.

## Notes

### Competing Interest Statement

The authors have declared no competing interest.

