## Supplementary figures and images for "Mutational Analysis of SARS-CoV-2 Genome in African Population"

### Supplementary file S2

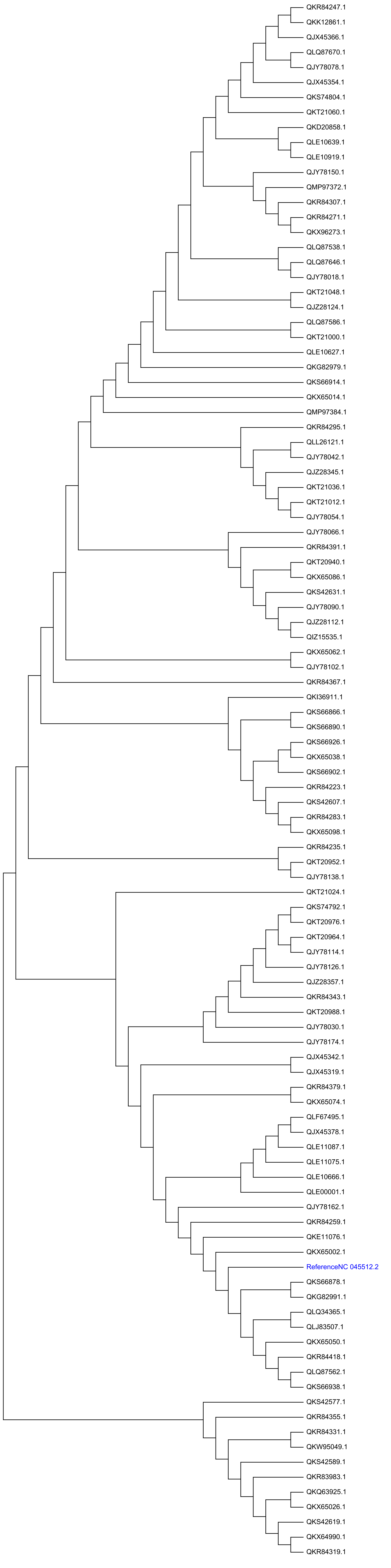
