## Supplementary file S1 for "Mutational Analysis of SARS-CoV-2 Genome in African Population": gisaid Uganda.pdf

We gratefully acknowledge the following Authors from the Originating laboratories responsible for obtaining the specimens, as well as the Submitting laboratories where the genome data were generated and shared via GISAID, on which this research is based.

All Submitters of data may be contacted directly via www.gisaid.org

| Accession ID | Originating Laboratory | Submitting Laboratory | Authors |
| --- | --- | --- | --- |
| EPI_ISL_451183, EPI_ISL_451185, EPI_ISL_451186, EPI_ISL_451190, EPI_ISL_451194, EPI_ISL_451198, EPI_ISL_451199, EPI_ISL_451200, EPI_ISL_451201, EPI_ISL_451202 | Uganda Virus Research Institute | MRC/UVRI & LSHTM Uganda Research Unit | Dan Lule Bugembe, John Kayiwa, My V.T Phan, Phionah Tushabe, Stephen Balinandi, Beatrice Dhaala, Deogratius Ssemwanga, Jonas Lexow, Henry Mwebesa, Jane Aceng, Henry Kyobe, Julius Lutwama, Pontiano Kaleebu, Matthew Cotten |
