## Supplementary file S1 for "Mutational Analysis of SARS-CoV-2 Genome in African Population": gisaid Zambia.pdf

We gratefully acknowledge the following Authors from the Originating laboratories responsible for obtaining the specimens, as well as the Submitting laboratories where the genome data were generated and shared via GISAID, on which this research is based.

All Submitters of data may be contacted directly via www.gisaid.org

| Accession ID | Originating Laboratory | Submitting Laboratory | Authors |
|---|---|---|---|
| EPI_ISL_510529 | School of Veterinary Medicine, Disease Control | School of Veterinary Medicine, Disease Control | Simulundu,E., Kapata,N., Mupeta,F., Kapata,P.C., Saasa,N., Changula,K., Muleya,W., Chitanga,S., Chambaro,H., Mubemba,B., Masahiro,K., Chanda,D., Mulenga,L., Fwoloshi,S., Shibemba,A.L., Kapaya,F., Zulu,P., Musonda,K., Monze,M., Sinyange,N., Liwewe,M.M., Kapin'a,M., Chipimo,P.J., Ngosa,W., Morales,A.N., Kayeyi,N., Malama,K., Tembo,J., Bates,M., Sawa,H., Takada,A., Nalubamba,K.S., Mukonka,V., Chilufya,C. and Zumla,A. |
